# fMRI-Based Prediction of Eye Gaze During Naturalistic Movie Viewing Reveals Eye-Movement–Related Brain Activity

**DOI:** 10.64898/2026.01.10.698820

**Authors:** Le Gao, Zhi Wei, Bharat B. Biswal, Xin Di

## Abstract

**Background:** Eye gaze provides crucial insights into perceptual and cognitive processes during naturalistic movie viewing, yet concurrent eye tracking is often unavailable in functional MRI (fMRI) research. While deep learning models can estimate gaze directly from fMRI eyeball signals, their out-of-the-box generalizability across heterogeneous datasets requires empirical evaluation.

**Methods:** We applied a specific pre-trained model from the DeepMReye framework in a zero-shot setting (without dataset-specific fine-tuning) to estimate gaze during movie watching across three independent fMRI datasets. Model accuracy was evaluated against camera-based eye-tracking data and via inter-subject correlations. Furthermore, we derived eye-movement-related time series from the predicted gaze signals to map their associated brain activation.

**Results:** At the individual level, predicted gaze showed modest correspondence with measured ground-truth data (*r* ≈ 0.24–0.37), yielding brain activation maps largely restricted to the visual cortex. In contrast, group-averaged gaze predictions exhibited substantially higher reliability (*r* ≈ 0.73–0.84). First-level General Linear Models (GLMs) derived from group-averaged predictions successfully revealed widespread activation across established oculomotor control regions, including the frontal and parietal eye fields. Exploratory analyses of age-related effects on gaze prediction and brain activity yielded inconsistent results across datasets.

**Conclusions:** Under a zero-shot implementation, the pre-trained model exhibits limitations for individual-level inference, likely reflecting the absence of dataset-specific training. However, group-averaged fMRI-based gaze estimates successfully capture shared viewing behaviors and robustly support the investigation of eye-movement-related brain activity. These findings inform the appropriate use of fMRI-based gaze decoding for naturalistic neuroimaging datasets lacking ground-truth eye-tracking logs.

## 1. Introduction

Eye movements are central to human perception and cognition. They guide visual exploration, support attentional selection, and facilitate social communication by revealing what people attend to and how they interpret dynamic visual scenes (Henderson 2003; Tatler et al. 2011). During naturalistic activities such as movie watching, gaze patterns reflect moment-to-moment fluctuations in engagement, comprehension, and social inference (Hasson et al. 2004; Richardson and Saxe 2020). Because of this richness, eye movements are widely used as behavioral markers for studying cognitive processes, including attention, memory, and theory-of-mind reasoning.

Functional MRI (fMRI) provides powerful tools for studying the neural mechanisms underlying these processes, but a major limitation in naturalistic fMRI research is that most datasets do not include simultaneous eye-tracking recordings. This gap is especially evident in large open-access datasets and legacy studies collected before eye-tracking systems became widely available (Sonkusare et al. 2019). Even today, many research sites lack in-scanner eye-tracking hardware, and when such systems are available, obtaining high-quality recordings is often difficult—particularly in populations such as young children, individuals with neurodevelopmental or psychiatric conditions, or participants who have difficulty maintaining stable head position. As a result, many potentially informative fMRI datasets lack reliable gaze information, complicating the interpretation of neural activity related to visual attention, scene processing, and oculomotor control and leaving valuable datasets under-utilized.

Recent advances in deep learning have introduced a possible solution. Models such as DeepMReye can infer gaze position directly from the MR signal of the eyeball region (Frey et al. 2021). These approaches build on earlier observations that motion of the eye and surrounding tissues produces systematic MR signal variations within the eyeball, likely reflecting contrast changes between internal ocular structures (Son et al. 2020). Such MR signals can be decoded using modern machine learning approaches, including convolutional neural networks. Subsequent work has further explored convolutional neural network-based approaches for MR-based gaze estimation (Wu et al. 2024), demonstrating the potential of deep architectures to capture gaze-related information from eyeball signals. The ability to estimate gaze retrospectively offers a powerful opportunity: large fMRI datasets without eye tracking can be “augmented” with reconstructed gaze trajectories, enabling novel analyses of visual behavior and attention without requiring camera-based systems.

Beyond methodological utility, fMRI-based gaze decoding opens new avenues for developmental and clinical research. Children and adults differ substantially in their visual exploration patterns, attentional control, and consistency of gaze behavior (Kirkorian et al. 2012). These behavioral differences correspond to ongoing maturation of large-scale cortical networks involved in visual attention, cognitive control, and social processing (Casey et al. 2005; Luna et al. 2015). Movie-watching paradigms are particularly well-suited for developmental studies because they elicit reliable neural responses across individuals while also capturing meaningful age-related differences in gaze patterns and comprehension (Cantlon and Li 2013; Petroni et al. 2018; Di et al. 2026).

In this study, we applied the pretrained DeepMReye to three naturalistic movie-watching fMRI datasets to assess its ability to reconstruct gaze behavior and to characterize the neural correlates of predicted eye movements. We first evaluated prediction accuracy by correlating model-based gaze estimates with simultaneously acquired camera-based eye-tracking data in a zero-shot setting. We also leveraged the well-established phenomenon of inter-individual correlation during shared video viewing (Hasson et al. 2004; Madsen and Parra 2022; Di et al. 2025), and quantified inter-individual correlations of the DeepMReye predictions themselves. If the model accurately captures gaze behavior, we expect both strong correlations with ground-truth eye tracking and high intersubject consistency. In addition, because gaze patterns show robust inter-individual correlations, we computed group-averaged gaze predictions and compared correlations with eye-tracking data at both the group and individual levels. This approach tests the hypothesis that even when individual predictions are noisy, averaging across participants may enhance signal quality and improve correspondence with ground-truth gaze.

Moreover, we examined brain activity associated with eye movements by using frame-to-frame gaze displacement (Euclidean distance) as a regressor in a conventional fMRI activation analysis. We hypothesized that we would identify regions involved in oculomotor control, such as the frontal eye fields (Pierrot-Deseilligny et al. 2004). We conducted these analyses using both group-averaged and individual-level gaze predictions to compare their ability to reveal eye-movement–related neural responses. Together, these analyses provide insight into the performance of the pretrained DeepMReye and its utility for studying gaze-related brain activity in naturalistic fMRI datasets.

## 2. Materials and Methods

An overview of datasets used and the analysis workflow is shown in Figure 1.

**Figure 1.**
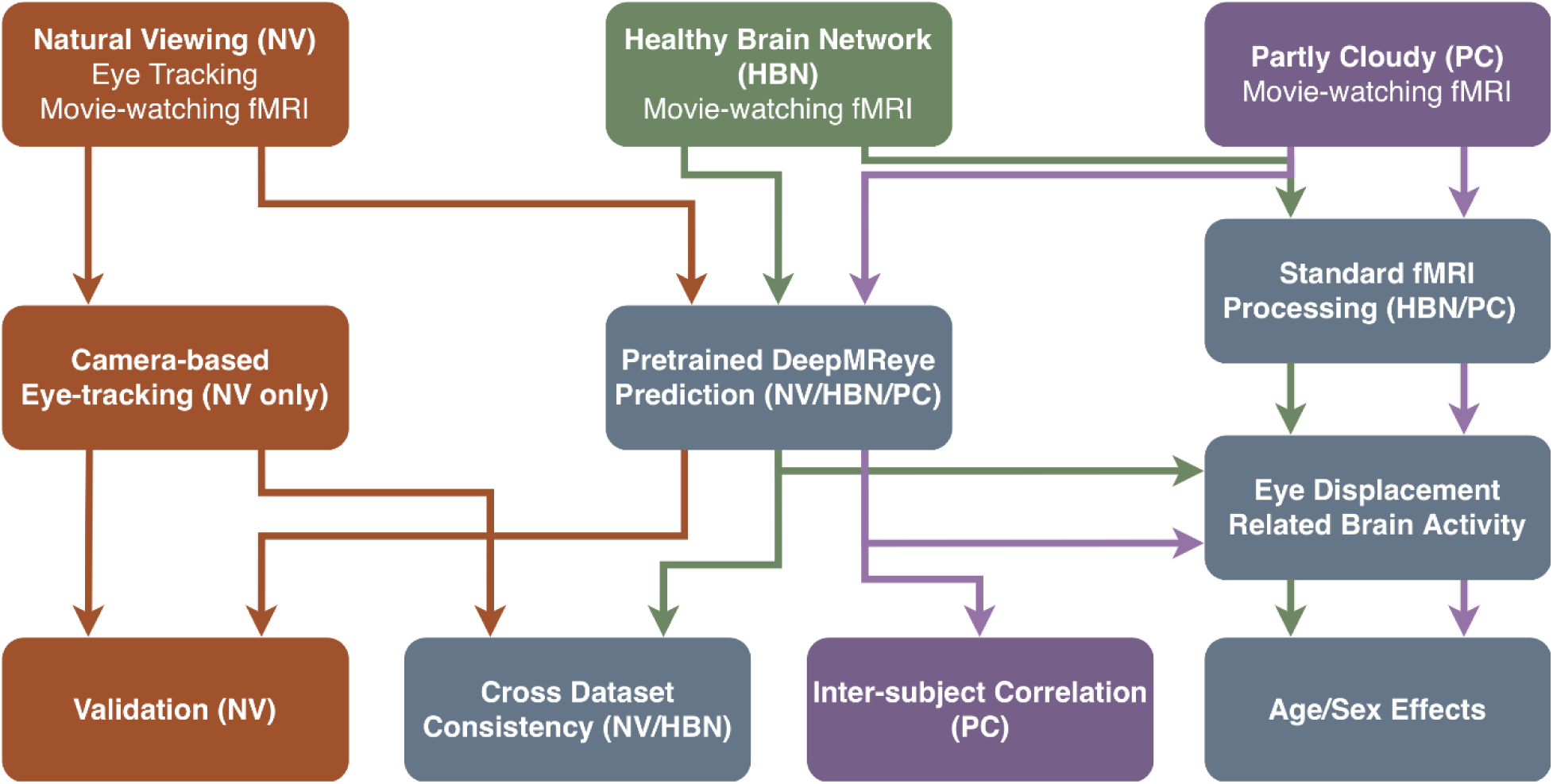
Study workflow for fMRI-based gaze decoding and downstream analyses. Eyeball fMRI voxels were extracted from three naturalistic viewing datasets—Natural Viewing (NV; brown), Healthy Brain Network (HBN; green), and Partly Cloudy (PC; purple)—following motion correction and template alignment. A pre-trained DeepMReye model then performed zero-shot inference to extract predicted gaze paths. These predictions underwent several levels of evaluation and downstream analysis: **(1) Validation:** NV predictions were compared against ground-truth camera-based eye-tracking data using correlation and Mean Absolute Error (MAE). **(2) Consistency:** Cross-dataset consistency was evaluated by correlating HBN predictions with NV ground-truth data for shared video stimuli, while inter-subject correlation (ISC) assessed cross-individual consistency in the independent PC dataset. **(3) Brain Mapping:** Gaze displacement (Euclidean distance) was calculated from the predicted time series and used as a regressor in a General Linear Model (GLM) to identify eye-movement-related brain activation. Finally, group-level analyses evaluated the effects of age and sex on these functional activation maps.

### 2.1. Participants and Datasets

This study utilized three publicly available fMRI datasets. All datasets included fMRI scans acquired during naturalistic movie-watching tasks, along with demographic information such as age and sex. Across all datasets, participants were included based on data quality criteria. Quality control procedures were applied at both the fMRI and eye-tracking levels. For the NV dataset specifically, eye-tracking data were evaluated post-preprocessing; rather than enforcing a rigid quantitative threshold, participants were excluded only if their recordings contained a critically insufficient amount of usable data.

**Natural Viewing (NV) Dataset** (Telesford et al. 2023): This dataset includes 22 adult participants (ages 23–51 years) and was collected at the Nathan S. Kline Institute. We focused on the naturalistic viewing portion where participants watched “The Present” (TP; 4min 18s) and a clip from “Despicable Me” (DM; 10min). The fMRI data was acquired with a repetition time (TR) of 2.1s. Critically, this dataset also contains simultaneously recorded eye-tracking data (EyeLink 1000 Plus, 250 Hz), which serves as a ground truth for validating the accuracy of the predicted gaze data from DeepMReye.

**Partly Cloudy (PC) Dataset** (Richardson and Saxe 2020): This dataset provided data from two distinct age groups, comprising 53 children (ages 3–12 years) and 29 adults (ages 18–39). All participants viewed a silent version of the animated short film “Partly Cloudy” (5min 36s). The fMRI scanning TR was 2.0s. This dataset does not include concurrent eye-tracking recordings.

**Healthy Brain Network (HBN) Dataset** (Alexander et al. 2017): This large-scale dataset includes data from children and young adults (ages 5–21 years). To avoid confounding factors from psychopathology, we included only healthy control participants. The stimuli were the same as in the NV dataset (TP and DM). The HBN data were acquired with a higher temporal resolution (TR = 0.8s), providing a more densely sampled BOLD signal. After applying quality control criteria, our final sample consisted of 82 participants for TP and 80 participants for DM. This dataset does not include concurrent eye-tracking recordings.

A summary of the three datasets is provided in **Table 1**. The datasets used in this study were acquired with different scanning protocols, including variations in voxel size, phase encoding direction, and scanner hardware. These differences may introduce domain shift between the training data of DeepMReye and the datasets analyzed here. More details about the study design and MRI acquisitions can be found in the respective papers.

**Table 1.** Summary of datasets used in this study.

| Dataset | N | Age Range (years) | Stimuli | TR (s) | Eye-tracking |
| --- | --- | --- | --- | --- | --- |
| NV | 22 | 23–51 | TP, DM | 2.1 | Yes |
| PC | 82 | 3–39 | PC | 2 | No |
| HBN | 82/80 | 5–21 | TP, DM | 0.8 | No |

*Note: TP = "The Present"; DM = "Despicable Me"; PC = "Partly Cloudy". For HBN, sample sizes differ between TP (n = 82) and DM (n = 80) due to data availability after quality control.*

### 2.2. fMRI Preprocessing

All fMRI data were preprocessed using SPM12 in MATLAB following a standardized pipeline (Di and Biswal 2023). The procedures for the PC and HBN datasets have been detailed previously (Di and Biswal 2022; Di et al. 2023); a brief summary is provided here. Structural images were segmented and normalized to MNI space. For *The Present* and *Despicable Me* datasets, the first 10 functional volumes were removed (yielding 240 and 740 volumes), whereas all volumes were retained for *Partly Cloudy*. Functional images were realigned, coregistered to the structural scan, normalized using the segmentation-derived deformation fields, and smoothed with an 8-mm FWHM Gaussian kernel.

### 2.3. Gaze Prediction from Pretrained DeepMReye

DeepMReye was applied in a zero-shot manner using the pretrained weights (datasets_1to6.h5) provided in the official DeepMReye repository without any fine-tuning on the target datasets. This design choice reflects the intended use case of applying pretrained DeepMReye to large-scale fMRI datasets that lack ground-truth eye-tracking data, where supervised fine-tuning is not feasible. Gaze coordinates (horizontal x, vertical y) were predicted from the fMRI time series using DeepMReye, a pre-trained deep neural network. All preprocessing steps followed the official DeepMReye pipeline. Eye voxels were extracted using the DeepMReye pipeline as described in Frey et al. (Frey et al. 2021), including non-linear alignment of the eye region to a template space followed by voxel selection based on the standardized eye mask provided with the model. Quality control reports generated by the DeepMReye pipeline were visually inspected for each dataset, and representative examples are provided in the Supplementary Materials. The model utilizes normalized eyeball voxel signals as input to estimate eye position in degrees of visual angle for each fMRI volume. The output was a time series of (x, y) gaze coordinates for each fMRI run.

### 2.4. Statistical Analysis of Gaze Movement

To quantitatively evaluate model performance, we assessed the correspondence between ground-truth eye-tracking data and predicted gaze movements using Pearson correlation analysis. For each participant and each viewing task, recorded and predicted gaze time series were temporally aligned. Individual correlation coefficients were computed separately for the horizontal (x) and vertical (y) gaze dimensions across all time points within a run. For group-level evaluations, we first computed group-averaged gaze trajectories by averaging the eye-tracking and predicted gaze time series across participants at each time point, yielding a summary time series for tracking and prediction, respectively. Pearson correlation coefficients were then calculated between the group-mean tracking and group-mean prediction trajectories for the x and y dimensions. This group-averaging approach emphasizes shared, stimulus-driven gaze dynamics by reducing idiosyncratic individual variability. In addition to Pearson correlation, we computed the Mean Absolute Error (MAE) for Natural Viewing Dataset between predicted and ground-truth gaze positions, expressed in degrees of visual angle. MAE quantifies the average spatial deviation and complements correlation by capturing absolute prediction error.

Eye-tracking data quality in the NV dataset was quantified using the proportion of valid samples after preprocessing and the root mean square (RMS) of sample-to-sample gaze displacement; their relationships with prediction accuracy are reported in Supplementary Figure. S2. On average, participants retained 79.4% (DM task) and 86.4% (TP task) of valid eye-tracking samples after preprocessing.

### 2.5. Statistical Analysis of fMRI Data

A two-level, voxel-wise general linear model (GLM) approach was employed to analyze the fMRI data in SPM12. At the first level, subject-specific effects related to gaze were modeled. At the second level, group-level effects and the relations to age and sex were assessed.

#### 2.5.1. First-Level Analysis: Subject-Specific GLM

For each participant, a first-level GLM was specified for each fMRI run to identify brain regions showing activity correlated with moment-to-moment changes in gaze. The design matrix for each participant included a primary regressor of interest and several nuisance regressors. The primary regressor was a parametric modulator representing gaze displacement (*d_t_*) calculated as the Euclidean distance between predicted gaze coordinates at successive time points:

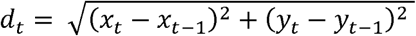

where (*x_t_, y_t_*) represents the predicted gaze position in degrees of visual angle at time point t. This displacement measure captures the magnitude of eye movement at each time point.

This displacement time series was then convolved with the canonical hemodynamic response function (HRF) to model the expected BOLD signal. To account for motion-related artifacts, 24 nuisance regressors were included in the model. These were derived from the 6 realignment parameters (3 translations and 3 rotations) obtained during preprocessing, their temporal derivatives, and quadratic terms of all 12 parameters. Following the model specification, the GLM was estimated for each subject. The full GLM equation for each voxel can be expressed as:

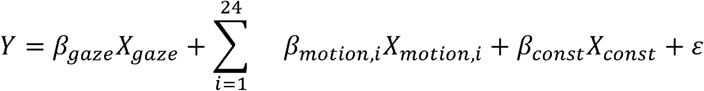

where Y is the BOLD signal at a given voxel, *β_gaze_* is the parameter estimate for the gaze displacement regressor, *β_motion,i_* represents the parameter estimate for the i-th motion regressor, *β_const_* is the intercept (mean signal level), and ε is the residual error term. This process generated a beta map for each participant, representing the effect size of the relationship between BOLD activity and gaze displacement.

#### 2.5.2. Second-Level Analysis: Group-Level Multiple Regression

To make inferences at the group level, the individual beta maps from the first-level analysis were entered into a second-level, whole-brain multiple regression model. This analysis aimed to identify brain regions where gaze-related activity was significantly associated with age and sex. The model included the following covariates: age (linear term), age² (quadratic term), and sex (coded as 0 = female, 1 = male).

For each voxel, the first-level beta value (B-) for subject j was modeled as:

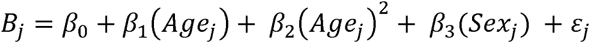

where β-is the intercept, β- and β-represent the linear and quadratic effects of age, respectively, β-represents the effect of sex, and ε-is the residual error for subject j. This model allowed us to examine both linear and non-linear relationships between age and brain activity, while controlling for sex.

### 2.6. Region of Interest (ROI) Analysis

To further characterize age-related effects identified in the whole-brain analysis, we conducted a Region of Interest (ROI) analysis. ROIs were defined based on clusters showing significant gaze-related activity (e.g., p < 0.001, FWE-corrected) from the group-level analysis, representing core regions of the eye-movement network. For each participant, we extracted the mean beta value from all voxels within each ROI from their first-level results. These values were then plotted against age to visualize the developmental trajectory of gaze-related activity.

## 3. Results

### 3.1. DeepMReye predictions vs. Eye-tracking in NV dataset

We first validated DeepMReye predictions using the NV dataset, which included concurrent camera-based eye-tracking data as ground truth. At the group level, the model performed well: group-averaged predicted gaze time series showed high correspondence with the group-averaged ground-truth eye-tracking data. As illustrated in **Figure 2a** and **d**, group-averaged predicted and actual gaze trajectories were closely aligned. For the DM stimulus, correlations were *r* = 0.84 (*p* < .001) horizontally and *r* = 0.73 (*p* < .001) vertically, and mean absolute errors were MAE = 0.564 horizontally and MAE = 0.486 vertically. Similarly, for TP stimulus, Pearson correlations were *r* = 0.81 (*p* < .001) for both horizontal and vertical gaze directions, and mean absolute errors were MAE = 0.550 horizontally and MAE = 0.380 vertically. These results demonstrate that DeepMReye reliably captures the group-level patterns during naturalistic viewing.

**Figure 2.**
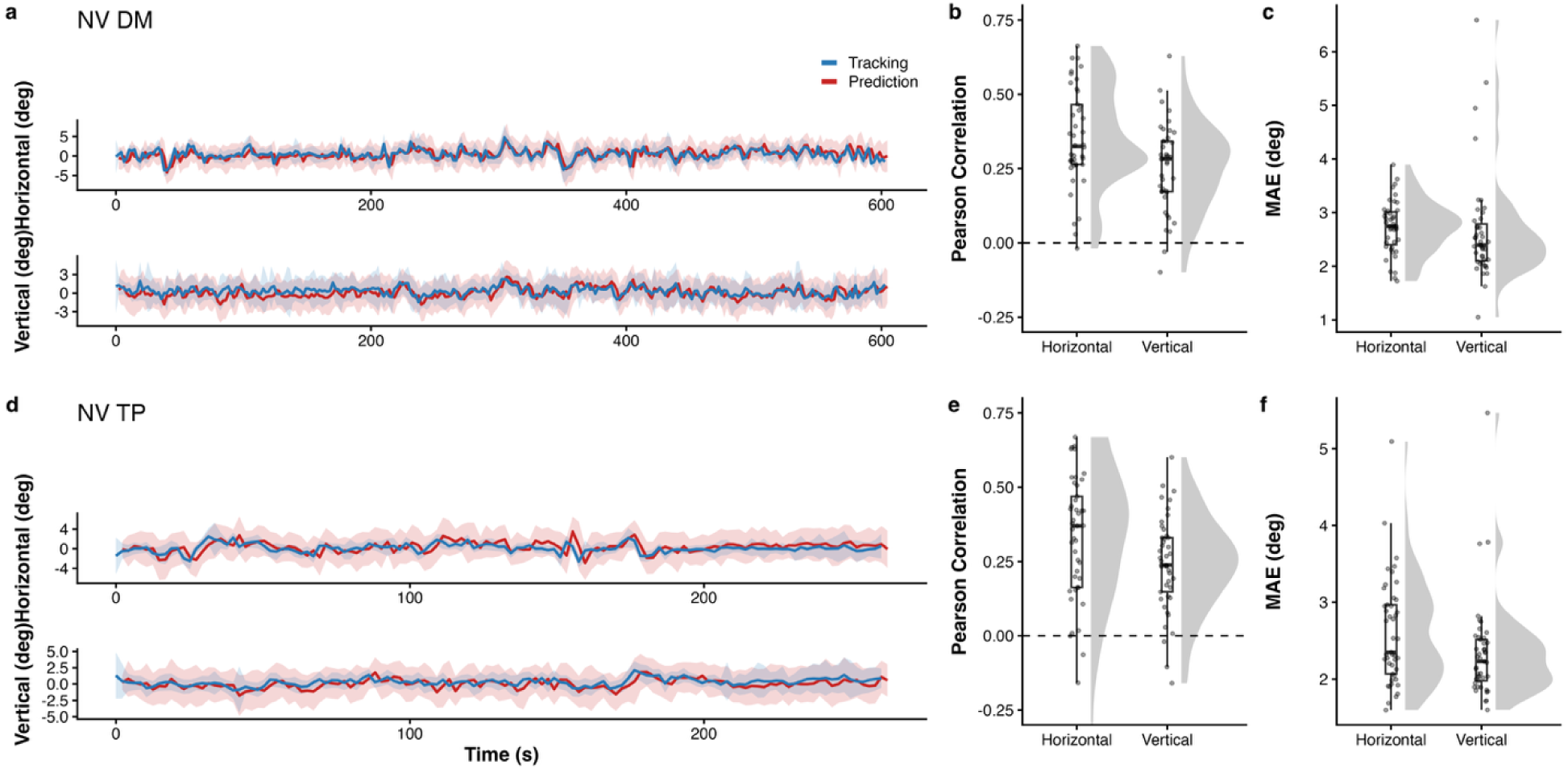
Validation of DeepMReye gaze predictions against ground-truth eye-tracking data. (a, d) Group-averaged horizontal and vertical gaze trajectories for the Natural Viewing (NV) dataset in the Despicable Me (DM) and The Present (TP) tasks. Solid lines represent the mean across subjects, and shaded regions indicate ±1 standard deviation. (b, e) Subject-level Pearson correlation coefficients between predicted and ground-truth gaze signals for horizontal and vertical components. (c, f) Subject-level mean absolute error (MAE) in degrees of visual angle, reported separately for horizontal and vertical components.

However, prediction accuracy declined substantially at the individual level. **Figure 2b** and **e** present raincloud plots of individual-participant Pearson correlation coefficients between predicted and ground-truth gaze for horizontal (x) and vertical (y) directions. Individual-level correlations showed greater variability and were generally lower than those observed at the group level. For the DM stimulus, median correlations were *r* = 0.32 (interquartile range [IQR] = 0.26–0.47) for horizontal and *r* = 0.28 (IQR = 0.17–0.34) for vertical gaze directions. For the TP stimulus, median correlations were *r* = 0.37 (IQR = 0.16–0.47) for horizontal and *r* = 0.24 (IQR = 0.15–0.33) for vertical gaze directions. While a subset of participants exhibited relatively high correlations between predicted and ground-truth gaze, many others showed substantially lower correspondence. Consistent with the correlation results, MAE values indicated moderate spatial accuracy at the individual level, with improved consistency observed after group averaging (**Figure 2c**, **f**). At the individual level, corresponding median spatial errors were MAE = 2.75° (IQR = 2.403– 3.011) horizontally and MAE = 2.40°(IQR = 2.101– 2.789) vertically for DM task, and MAE = 2.35° (IQR = 2.068– 2.968) horizontally and MAE = 2.23°(IQR = 1.978– 2.514) vertically for TP task. Together, these results indicate that individual-level prediction performance is more variable and generally weaker than group-level performance.

### 3.2. Cross-Dataset Consistency between NV and HBN datasets

To further assess the generalizability of pretrained DeepMReye, we performed a cross-dataset evaluation by comparing group-averaged predictions from the HBN dataset with ground-truth eye-tracking data from the NV dataset. Cross-dataset analyses were performed to assess the consistency of group-level gaze dynamics across datasets, rather than to provide a direct measure of prediction accuracy. Both datasets used identical stimuli (TP and DM), allowing direct comparison despite differences in participant samples and scanning parameters. As shown in **Figure 3a** and **d**, the predicted gaze trajectories from HBN participants closely matched the ground-truth data from NV participants. For the DM stimulus, Pearson correlations were *r* = 0.87 for horizontal and *r* = 0.80 for vertical gaze directions. For the TP stimulus, correlations were *r* = 0.90 for horizontal and *r* = 0.86 for vertical gaze directions. These results indicate strong correspondence in group-level, stimulus-driven gaze dynamics across datasets.

**Figure 3.**
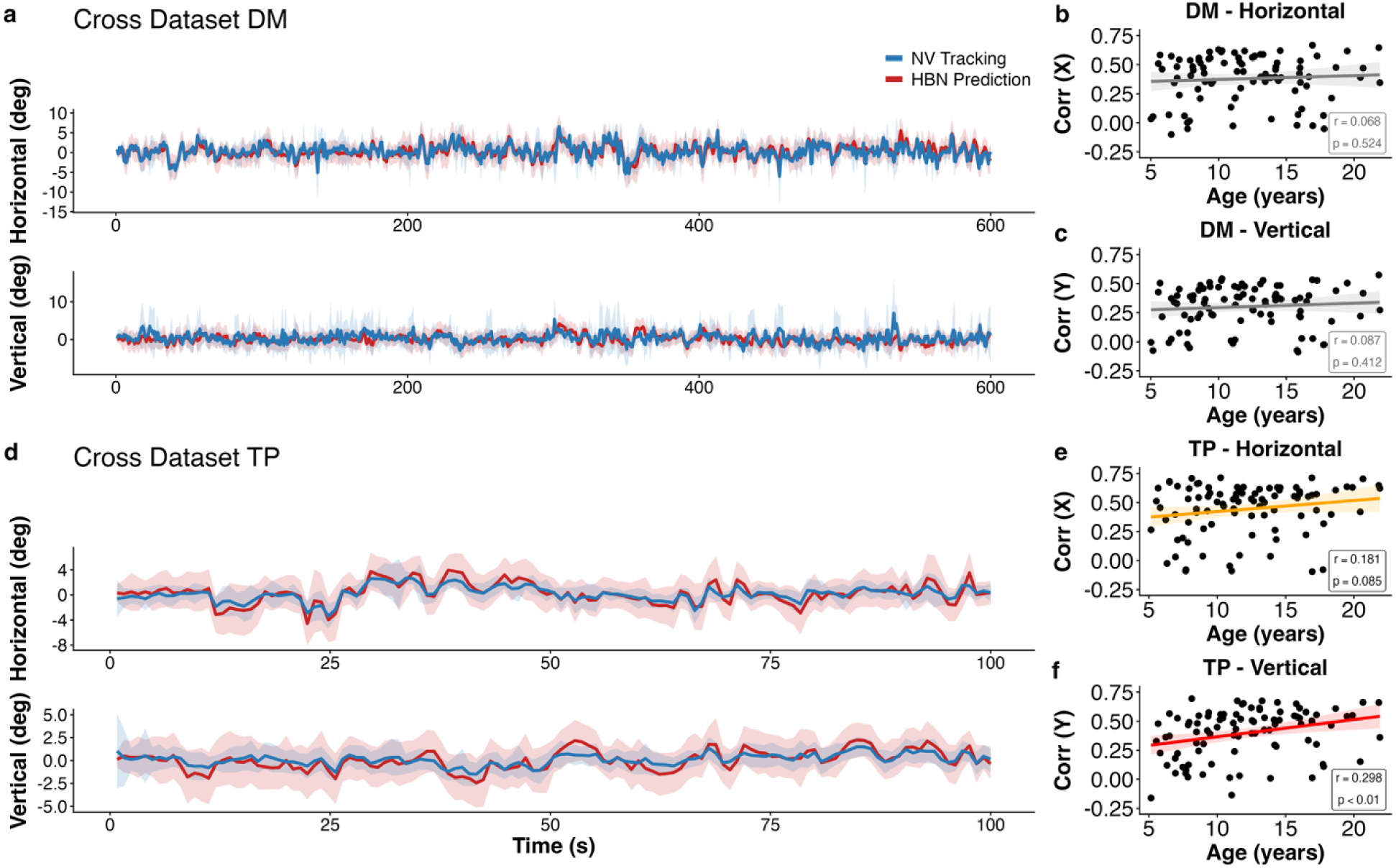
Cross-dataset comparison of group-mean gaze trajectories and age-related correlation patterns between the ground-truth of the Natural Viewing (NV) dataset and the predicted gaze of the Healthy Brain Network (HBN) dataset. (a, d) Group-averaged gaze trajectories for the cross-dataset comparison between NV ground-truth eye tracking (blue) and HBN predicted gaze signals (red) in the Despicable Me (DM) and The Present (TP) tasks, respectively. Solid lines represent the mean across subjects within each dataset, and shaded regions indicate ±1 standard deviation. Horizontal and vertical gaze components are shown separately. (b, c) Associations between age and subject-level similarity to the NV group-average gaze trajectory in the HBN dataset for the DM stimulus, shown separately for horizontal and vertical components. Each point represents one HBN subject/run, and the fitted line indicates the linear trend. (e, f) Associations between age and subject-level similarity to the NV group-average gaze trajectory in the HBN dataset for the TP stimulus, shown separately for horizontal and vertical components.

We further quantified this relationship by computing the correlation between each participant’s predicted gaze trajectory and the NV group-mean tracking gaze trajectory. During the DM task, positive associations were observed for both horizontal (r = 0.068, p = 0.524) and vertical (r = 0.087, p = 0.412) gaze components, though these effects did not reach statistical significance (**Figure 3b** and **c**). In contrast, during the TP task, stronger age-related increases in correlation were observed, particularly for the vertical gaze dimension (r = 0.298, p = 0.004), with a weaker trend for the horizontal component (r = 0.181, p = 0.085) (**Figure 3e** and **f**).

These results indicate that individual gaze predictions increasingly align with normative group-level viewing patterns as age increases, especially under structured viewing conditions. This finding suggests improved model generalization and developmental stabilization of gaze behavior in later childhood and adolescence.

### 3.3. Intersubject correlations of predicted gaze in PC dataset

To further examine developmental effects on gaze behavior, we analyzed intersubject similarity and variability in the PC dataset (**Figure 4**). Correlation matrices revealed increasingly structured alignment between the same group of individual predictions with increasing age (**Figure 4a** and **b**). The distinct high-correlation cluster among adults in the lower-right quadrant of the matrices, together with the weaker correlations involving younger participants, highlights age-related differences in gaze synchrony during movie viewing.

**Figure 4.**
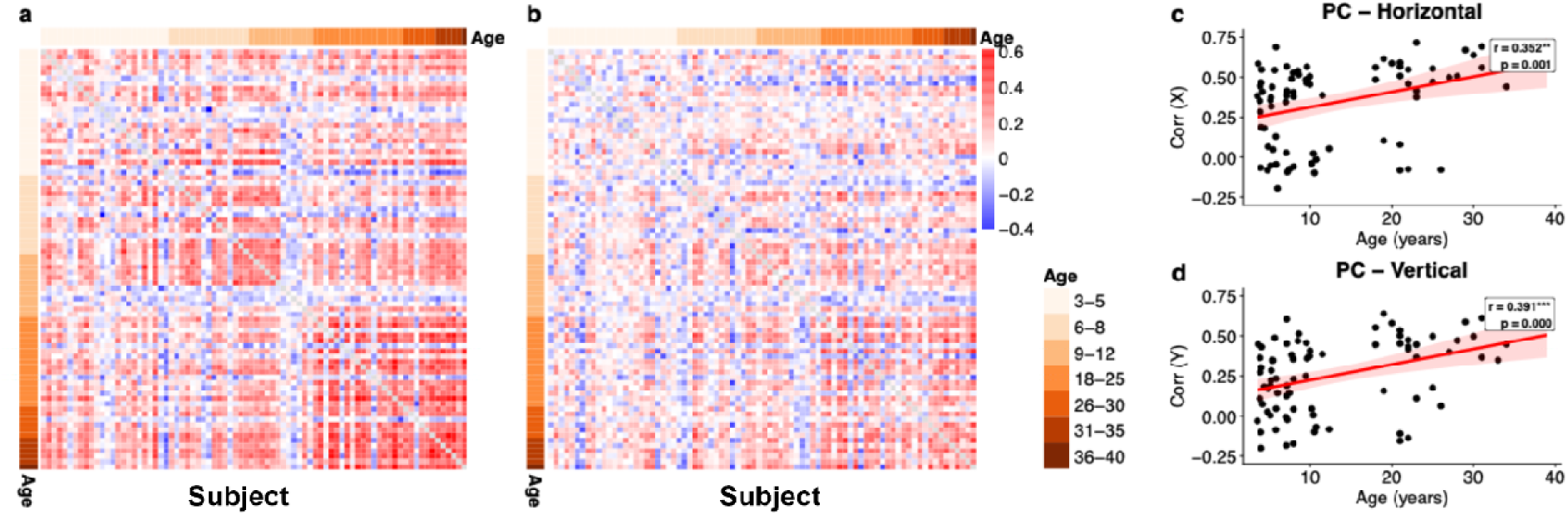
Intersubject similarity and variability of gaze patterns in the Partly Cloudy dataset. (a–b) Intersubject correlation matrices within the PC dataset revealed increasingly structured alignment of predicted gaze trajectories with increasing age for horizontal (a) and vertical (b) gaze components. (c–d) Scatter plots showing the relationship between age and correlation strength for horizontal (c) and vertical (d) gaze components. Solid lines indicate linear regression fits, and shaded regions represent 95% confidence intervals. Red lines indicate statistically significant age effects (*p* < 0.05).

Quantitative analyses demonstrated a significant positive relationship between age and gaze similarity for both horizontal (r = 0.352, p = 0.001) and vertical (r = 0.391, p < 0.001) components (**Figure 4c** and **d**), indicating that older participants exhibited stronger correspondence with normative gaze patterns. These findings suggest age-related increases in gaze synchrony, potentially reflecting maturation of visual exploration strategies and improved coordination between attentional control and eye movement dynamics.

### 3.4. Brain activations Associated with Eye Movements

To identify the core neural network associated with eye movements during naturalistic movie viewing, we construct eye-movement time series by calculating the Euclidean distance between gaze positions between consecutive time points, using either individual-level predicted gaze or group-averaged gaze predictions. Group-level analyses were performed separately for each movie.

As illustrated in **Figure 5a** and **c**, group-averaged eye-movement regressors produced highly similar whole-brain activation patterns for the HBN dataset during the movie-watching task DM and for the PC dataset during movie-viewing task. Robust activation was consistently observed in a distributed oculomotor control network, with prominent involvement of frontal and parietal regions implicated in the planning and control of eye movements, including the frontal eye fields (FEF) and the intraparietal sulcus (IPS). These regions constitute core components of the dorsal attention and oculomotor networks that support saccade generation, visuospatial attention, and top-down control of visual exploration.

**Figure 5.**
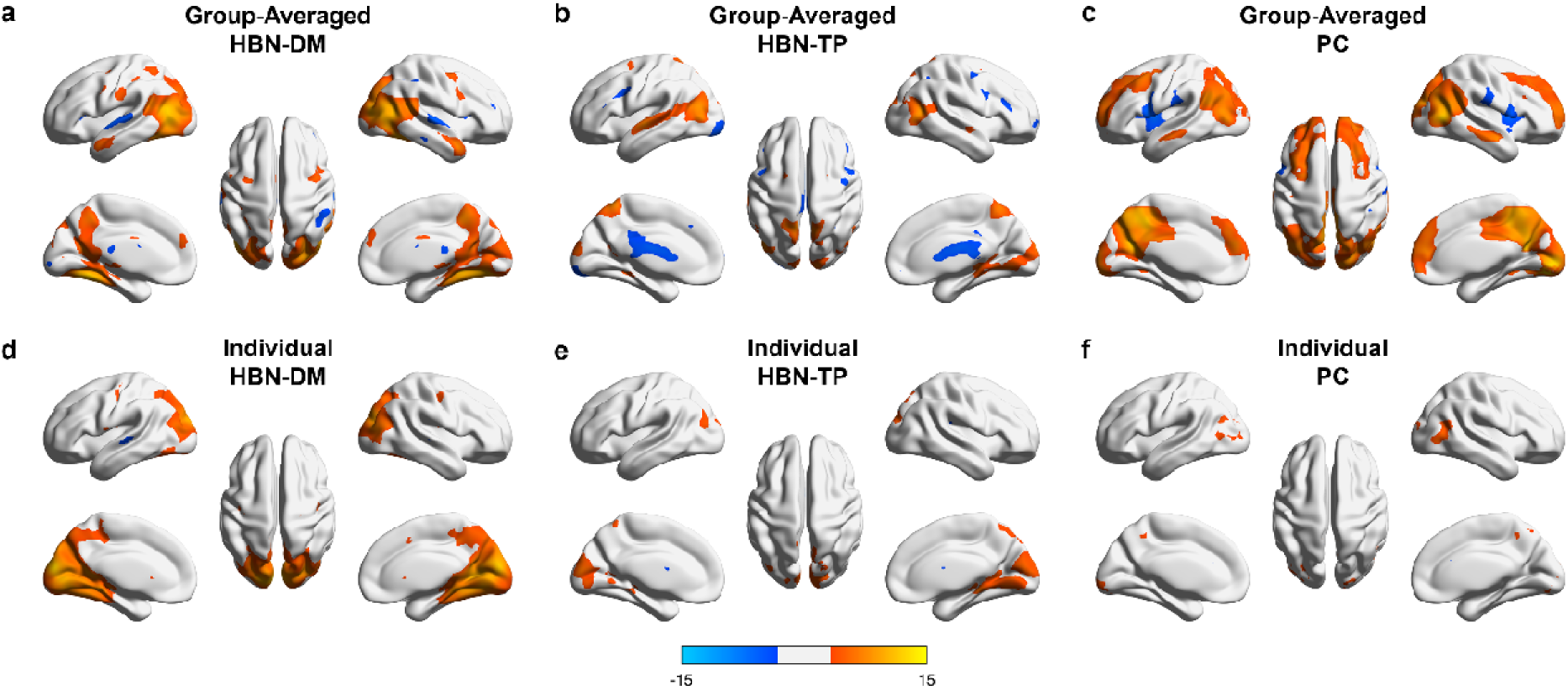
Main Effect of Gaze Displacement on Whole-Brain Activation. The figure shows regions where brain activity was significantly and positively correlated with individual-level and group-averaged gaze displacement. Activation using group-averaged gaze prediction of (a) *Despicable Me* (DM) stimulus of the Healthy Brain Network (HBN) dataset, (b) *The Present* (TP) stimulus of the HBN dataset, and (c) Partly Cloudy (PC) dataset. Activation using individual gaze prediction of (d) DM stimulus of the HBN dataset, (e) TP stimulus of the HBN dataset, and (f) PC dataset. Warm colors indicate a stronger positive correlation, and cold colors indicate negative correlations.

In addition to this control network, widespread activation was observed in occipital visual cortices, encompassing the cuneus, lingual gyrus, and primary visual cortex, reflecting the expected engagement of early visual processing during continuous gaze shifts. Notably, despite substantial differences between the HBN and PC datasets—including movie stimuli, participant age distributions, and fMRI acquisition parameters—the oculomotor control network associated with gaze displacement was largely conserved across datasets. This robust main effect provides a consistent basis for subsequent analyses examining how age modulates gaze-related brain activity. Compared with the DM movie-watching task, the TP task in the HBN dataset exhibited fewer and more spatially restricted activation clusters at the same statistical threshold (**Figure 5b**), potentially reflecting reduced statistical power due to shorter task duration and more constrained viewing dynamics.

In contrast, regressors derived from individual-level eye-movement predictions resulted in significantly smaller and more localized activation clusters across all three conditions. As illustrated in **Figures 5d** and **5e**, the DM and TP movie-watching tasks in the HBN dataset still showed positive activation in visual cortical regions (e.g., lingual gyrus and cuneus), but these effects were substantially less extensive. The PC task showed minimal and sparse activation at the same statistical threshold (**Figure 5f**). Overall, individual-level regressors yielded less extensive and less consistent group-level activation patterns compared to group-averaged regressors.

### 3.5. Age Effects on Eye-Movement related Brain activations

We further examined whether eye-movement–related brain activity was modulated by age by including age as a covariate in the second-level analyses. Age effects were evaluated separately for each dataset and for regressors derived from group-averaged versus individual-level gaze predictions. We found that the relationship between age and brain activity demonstrated different patterns across the two datasets, which featured different movie stimuli and participant cohorts. Similarly, we also compared the activation results derived from the beta values generated based on individual and group mean predictions.

#### 3.5.1. Age Effects in the Partly Cloudy (PC) Dataset

In the PC dataset, group-level analyses using group-averaged gaze predictions revealed significant positive age-related effects in the Postcentral Gyrus, the Precuneus, and the Middle Frontal Gyrus (**Figure 6a**). This pattern indicates systematic age-related modulation of gaze-related neural activity in regions implicated in sensory integration, visuospatial processing, and higher-order cognitive control during viewing of this stimulus. In contrast, no significant age-related activation was detected in analyses based on individual-level gaze predictions. When individual-level gaze predictions were used, age-related effects were weaker and spatially more restricted, highlighting the greater robustness of group-averaged gaze estimates for detecting developmental effects.

**Figure 6.**
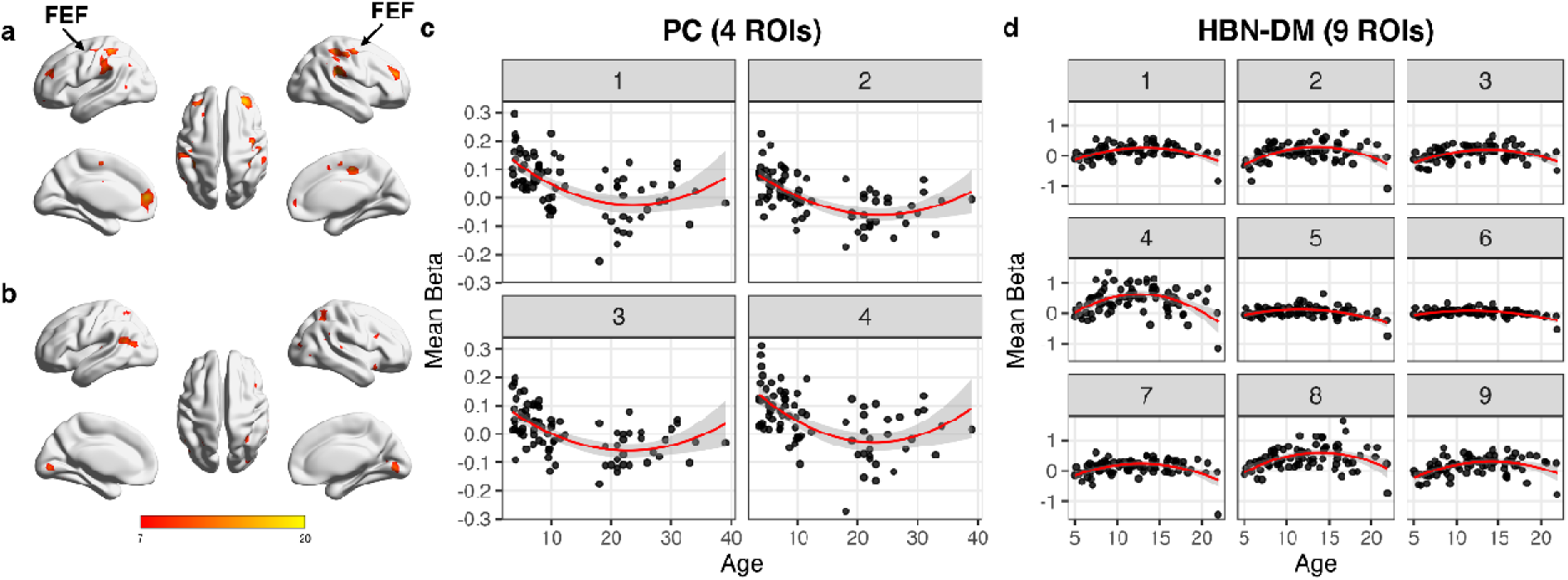
Modulation of Gaze-Related Brain Activity by Age. (a-b) Regions where brain activity exhibited a significant positive correlation with age using group-averaged gaze prediction from the Partly Cloudy (PC) dataset and *Despicable Me* (DM) stimulus of the Healthy Brain Network (HBN) dataset. (c-d) Scatter plots of mean beta values are versus age in significant ROIs based on results from the PC dataset and the DM stimulus of HBN dataset. Solid lines depict ROI-wise quadratic fits estimated using generalized linear models (Mean Beta ∼ Age + Age²), with shaded areas indicating 95% confidence intervals.

#### 3.5.2. Age Effects in the Healthy Brain Network (HBN) Dataset

The DM task in the HBN dataset, which involved viewing a clip from movie DM with complex dialogue and social scenes, revealed a distributed pattern of age-related effects based on group-averaged gaze predictions (Figure **6b**). Significant positive age-related associations were observed primarily in posterior visual regions, including the lingual gyrus and cuneus, as well as in the middle frontal gyrus. These regions are implicated in visual processing, visuospatial integration, and higher-order cognitive control, suggesting age-related modulation of gaze-related neural activity during naturalistic viewing. In contrast, no significant age-related effects were detected for the TP task.

### 3.6. ROI analysis

To assess age-related effects within these regions, we performed ROI-based analyses using quadratic regression models, with mean beta values extracted from each ROI as the dependent variable and both linear (Age) and quadratic (Age²) terms included as predictors (Mean Beta ∼ Age + Age²). Models were fitted separately for each ROI to characterize potential non-linear developmental trajectories.

In the PC dataset, quadratic modeling revealed significant age-related curvature in several ROIs, as indicated by a significant quadratic (Age²) term (Figure 5c). Specifically, gaze-related activation within these regions exhibited a non-linear developmental trajectory, characterized by higher activation in younger participants followed by a gradual decrease with increasing age. This pattern suggests that age-related changes in gaze-related brain activity in the PC dataset are not well described by a simple linear trend but instead reflect systematic non-linear modulation across development. In the HBN DM dataset, ROI-wise quadratic models likewise revealed significant quadratic age effects in multiple ROIs (Figure 5d). In contrast to the PC dataset, these regions showed inverted U-shaped relationships, with gaze-related activation increasing from childhood into adolescence and subsequently decreasing into young adulthood. The presence of significant quadratic terms indicates that age-related modulation of gaze-related neural activity during the DM task varies across developmental stages rather than progressing monotonically with age. Taken together, these ROI-based analyses demonstrate that age-related changes in gaze-related brain activity are non-linear and context-dependent, differing across datasets and stimulus conditions.

Whereas the PC dataset is characterized primarily by a decreasing trajectory with age, the HBN DM dataset exhibits developmentally specific peak activation during adolescence, underscoring the importance of modeling non-linear age effects when examining gaze-related neural development.

## 4. Discussion

In this study, we demonstrate that although a pretrained DeepMReye model yields limited accuracy at the individual level, its predictions become highly reliable when aggregated across participants, providing a potential approach for extracting gaze information from large-scale naturalistic fMRI datasets lacking eye-tracking. Although fine-tuning has been shown to improve performance when ground-truth eye-tracking data are available, our zero-shot evaluation highlights the extent to which a pretrained model from the DeepMReye framework can generalize across datasets without dataset-specific adaptation. At the behavioral level, model-derived predictions closely matched ground-truth eye-tracking data at the group level, both within and across datasets, while individual-level prediction accuracy was more variable. At the neural level, gaze-related activity consistently engaged a canonical oculomotor control network encompassing frontal and parietal regions, including the frontal eye fields and intraparietal sulcus, alongside expected activation in visual cortices. Importantly, age-related modulation of gaze-related brain activity differed across datasets, with non-linear developmental trajectories observed in ROI-based analyses.

As established in our analyses of the NV dataset, the group-averaged predictions demonstrated substantially higher fidelity to the ground-truth data. By averaging across participants, idiosyncratic variations and model prediction noise are attenuated, thereby enhancing the shared, stimulus-driven component of the gaze signal. This improvement in signal-to-noise ratio was directly mirrored in our neuroimaging results: brain activation maps derived from the group-mean regressor revealed a comprehensive and neuroanatomically plausible network, whereas maps from individual regressors yielded more restricted and variable activations.

The strong performance observed at the group level likely reflects the dominant influence of shared, stimulus-driven gaze patterns in naturalistic viewing. Prior work has demonstrated that movies strongly synchronize attention and gaze across viewers, producing robust inter-subject correlations in both behavior and neural responses (Hasson et al. 2004; Dorr et al. 2010; Mital et al. 2011). By averaging predicted gaze across participants, idiosyncratic noise arising from anatomical variability, voxel-level signal heterogeneity in the orbit, and motion-related artifacts is attenuated, yielding a clearer estimate of the shared component of gaze behavior. These findings highlight a key strength of applying pretrained MR-based gaze decoding models in naturalistic paradigms: despite the relatively coarse spatiotemporal resolution of fMRI, the shared component of visual attention elicited by naturalistic stimuli can be reliably recovered under this implementation.

In contrast, predictive correspondence at the individual level was markedly lower and more variable. Multiple factors may contribute to this limitation, including inter-individual differences in eye anatomy, orbit geometry, EPI distortions, and head motion, all of which can affect the extraction of reliable features from signals surrounding the eyeballs (Petit and Haxby 1999; Beauchamp 2003). In addition, because the pre-trained model was not fine-tuned on our datasets, differences in scanner characteristics, acquisition parameters, and population demographics may reduce generalizability (Tran et al. 2025; Botvinik-Nezer et al. 2020). Further, the temporal resolution of fMRI constrains the detection of rapid eye movements, making it difficult to capture saccades or subject-specific patterns of visual exploration (Engbert and Kliegl 2003). Together, these factors suggest that while the model captures shared, stimulus-driven gaze components, the reduced individual-level performance observed here likely reflects the absence of dataset-specific training and domain adaptation, rather than an intrinsic limitation of the DeepMReye framework itself. It is further supported by the weak or absent eye-movement–related activations observed when individual-level predictions were used. Eye-tracking data quality metrics were not consistently associated with prediction accuracy across tasks (Supplementary Figure. S2), suggesting that measurement noise alone does not fully account for the observed variability in individual-level performance. In addition, the relationship between predicted error (PE) and decoding performance was not consistent across tasks or evaluation metrics (Supplementary Figure. S3), indicating that PE should be interpreted cautiously as a proxy for model reliability in this zero-shot setting. Specifically, PE showed associations with MAE but not correlation in the TP task, whereas in the DM task the pattern differed, indicating task-dependent relationships between PE and performance metrics. These findings further suggest that factors such as domain shift and lack of dataset-specific adaptation may play a more prominent role in limiting individual-level performance. Importantly, the consistency of group-level predictions observed across datasets further suggests that the pretrained model captures robust and generalizable components of gaze behavior. Despite the aforementioned limitations and factors, the robustness to domain shift at the group level contrasts with the variability observed at the individual level, where subject-specific anatomical and noise-related factors play a larger role. Together, these findings indicate that while zero-shot models may be limited for individualized inference, they retain substantial utility for extracting reliable, population-level gaze signals across heterogeneous datasets.

These findings have important methodological implications for fMRI studies that lack camera-based eye tracking. Group-averaged gaze predictions can serve as a practical substitute for hardware-based eye tracking in studies focused on population-level gaze dynamics or stimulus-driven attentional processes, particularly in large naturalistic datasets where eye tracking was not acquired (e.g., HCP, NSD), when using pretrained models in a zero-shot setting. However, the current approach is not suitable for applications requiring precise individual-level gaze measurement, such as quality control of fixation compliance, fine-grained behavioral analyses, or clinical assessments at the single-subject level, where camera-based eye tracking remains essential.

It is important to acknowledge the inherent temporal limitations of fMRI when interpreting these oculomotor brain maps (the ’Nyquist problem’). Because the TRs in these datasets (800–2100 ms) are substantially longer than typical saccade durations (20–200 ms), the gaze signal is temporally aliased, making our ’gaze displacement’ metric mathematically unable to distinguish between one large saccadic shift and several smaller ones within a single volume. However, because the BOLD signal is inherently slow and acts as a low-pass filter, the high-frequency temporal precision of camera-based eye tracking is largely lost when convolved with the hemodynamic response function. Therefore, activation maps derived from fMRI-based gaze displacement are unlikely to differ substantially from those derived from high-resolution eye tracking, as both ultimately capture the same slow, aggregate oculomotor variance and macroscopic shifts in visual attention.

A central neuroscientific finding of this study is that age-related modulation of gaze-associated brain activity is non-linear and highly context dependent. In the HBN dataset, which spanned childhood through young adulthood and involved viewing a socially complex movie clip (DM), ROI analyses inverted U-shaped developmental trajectories in multiple regions within a distributed posterior–frontal network. These trajectories were characterized by increasing gaze-related activation from childhood into adolescence, followed by a subsequent decrease into young adulthood. Such patterns are consistent with prior work highlighting non-linear maturation of attentional and visual systems across development (Madden 2007). In contrast, the PC dataset exhibited a different pattern of age-related modulation, with gaze-related activity showing a predominantly decreasing trajectory across the sampled age range. Given the limited adolescent coverage in this dataset, these effects likely reflect differences between age groups rather than a fully resolved continuous developmental trajectory. Together, these findings emphasize that apparent age effects in naturalistic viewing are strongly shaped by both stimulus properties and sampling structure and are not adequately captured by simple linear models.

The presence of age-related differences in group-level analyses primarily suggests that the model may be sensitive to broad developmental trends in gaze behavior (Fernandes et al. 2024), but insufficiently precise to detect subject-specific effects amid individual noise. Whether these age effects reflect true changes in visual attention or methodological artifacts remains unclear. Future work should examine how demographic variables influence model performance and whether targeted model adaptation can improve sensitivity to individual differences.

Our findings revealed a notable dissociation between horizontal and vertical eye movement synchronization across development. While horizontal eye movement synchrony remained relatively stable across age groups, vertical eye movement synchrony showed a significant positive correlation with age (r = 0.298, p = 0.004). This differential developmental trajectory may reflect the distinct neural substrates underlying horizontal and vertical saccade generation. The prolonged maturation of vertical eye movement control suggested by these findings aligns with previous literature suggesting that certain oculomotor functions continue to develop throughout childhood and adolescence (Grönqvist et al. 2006).

These results indicate that age-related differences in gaze-related brain activity are strongly shaped jointly by the developmental stage and the stimulus context. Group-averaged model-derived gaze predictions are sufficiently robust to reveal population-level developmental patterns, whereas individual-level predictions remain too noisy to support reliable subject-specific inference. This highlights an important practical boundary of current zero-shot applications of pretrained fMRI-based gaze decoding models: deep-learning–based MRI gaze decoding may be valuable for identifying population-level developmental trajectories, but its current accuracy limitations constrain its utility for fine-grained individual differences work.

Several limitations should be noted. Our analyses were based on a limited number of movie stimuli and datasets, with camera-based eye tracking available only for one dataset. The use of a pre-trained model without dataset-specific fine-tuning may have constrained performance particularly in populations that differ substantially from the training sample. In addition, the temporal resolution of fMRI inherently limits the model’s sensitivity to rapid eye-movement events. These limitations motivate further methodological innovation.

Taken together, our findings should be interpreted in the context of a zero-shot application of a pretrained model. While this approach provides a practical solution for large datasets lacking eye tracking, it does not fully reflect the capabilities of the DeepMReye framework when combined with dataset-specific fine-tuning, standardized preprocessing pipelines, and rigorous quality control. Thus, the current results should be interpreted as a lower-bound estimate of performance under zero-shot conditions. Future work incorporating these elements will be essential to determine the full potential of MR-based gaze decoding for individual-level inference.

Recent work (Park et al. 2025) has highlighted the influence of motion artifacts and preprocessing strategies on the performance of DeepMReye, demonstrating that improved correction methods can enhance decoding accuracy. Our study focuses on a complementary use case, namely the application of DeepMReye in a zero-shot manner across heterogeneous naturalistic datasets, where advanced motion correction or fine-tuning may not be feasible. In this context, the robust group-level consistency observed here suggests that shared gaze-related signals can still be reliably extracted despite these challenges.

Future work should explore strategies for improving individual-level prediction accuracy, including incorporating subject-specific calibration (Kassner et al. 2014), retraining or fine-tuning models on larger or more diverse datasets (He et al. 2016), integrating high-resolution structural data of the orbit, and developing hybrid models that jointly use anatomical and functional information. Future work should also explore extending fMRI-based gaze decoding to predict discrete gaze events (e.g., fixations and saccades) rather than continuous position estimates, as well as integrating multimodal signals such as pupillometry or eyelid dynamics. Such improvements will be essential for enabling individualized inference and expanding the applicability of MR-based gaze decoding within the DeepMReye framework and related approaches, particularly for studying individual differences in clinical populations such as autism spectrum disorder (ASD) or attention-deficit/hyperactivity disorder (ADHD), where gaze behavior represents a key behavioral marker.

## Conclusion

In summary, this study demonstrates that fMRI-based gaze prediction provides a potentially useful framework for investigating gaze-related brain activity at the group level during naturalistic viewing, while also highlighting current limitations for reliable individual-level inference. By validating a pretrained DeepMReye model in the Natural Viewing dataset using concurrent camera-based eye-tracking data, and by evaluating its zero-shot performance through inter-subject correlation and downstream analyses, we show that model-derived gaze estimates recover biologically meaningful oculomotor and attentional networks and reveal systematic, age-related modulation of gaze-related brain activity.

Importantly, these developmental effects are not uniform across contexts but vary as a function of stimulus properties, task demands, and population characteristics, underscoring the context-dependent nature of neurodevelopment during naturalistic viewing. Together, these findings highlight the value of integrating data-driven computational approaches with large-scale developmental neuroimaging datasets to move beyond static, task-isolated paradigms, offering a promising path toward understanding how dynamic visual behavior and its neural substrates evolve across development, particularly in settings where traditional eye tracking is unavailable.

## Data and Code Availability

MRI and eye-tracking data were obtained from publicly available sources: the Natural Viewing (NV) dataset (https://fcon_1000.projects.nitrc.org/indi/retro/nat_view.html), the Healthy Brain Network (HBN) dataset (http://fcon_1000.projects.nitrc.org/indi/cmi_healthy_brain_network/), and the OpenNeuro repository (https://openneuro.org/datasets/ds000228). Analysis scripts will be shared on OSF.

## Author Contributions

L.G. designed the study, implemented the gaze prediction, performed the analyses, and drafted the manuscript. X.D. provided overall conceptual guidance and contributed to the study design. B.B.B and Z.W. offered critical feedback and revisions throughout the project. B.B.B. and X.D. provided resources and secured funding for the project. All authors contributed to the interpretation of results and approved the final version of the manuscript.

