## Supplementary Materials for "fMRI-Based Prediction of Eye Gaze During Naturalistic Movie Viewing Reveals Eye-Movement–Related Brain Activity"

The supplementary materials provide additional methodological validation and quality control (QC) analyses supporting the main manuscript. First, to confirm the integrity of our preprocessing pipeline, we present representative QC reports demonstrating successful eye-voxel extraction and alignment across all three datasets (Figure S1). Second, to address the potential influence of ground-truth measurement noise on individual-level decoding accuracy, we quantify the relationship between camera-based eye-tracking data quality (data retention and spatial precision) and model prediction accuracy in the Natural Viewing dataset (Figure S2). Finally, we assess the utility of the model's unsupervised Predicted Error (PE) metric by examining its relationship to both spatial accuracy (Mean Absolute Error) and correlational performance (Figure S3), further characterizing the boundaries of individual-level gaze decoding.

**Supplementary Figure S1. Quality control of eye-voxel extraction across datasets.** Representative quality control (QC) visual reports generated by the DeepMReye preprocessing pipeline. Panels display the successful alignment and extraction of eye voxels for a representative participant from the (a) Natural Viewing (NV), (b) Healthy Brain Network (HBN), and (c) Partly Cloudy (PC) datasets.

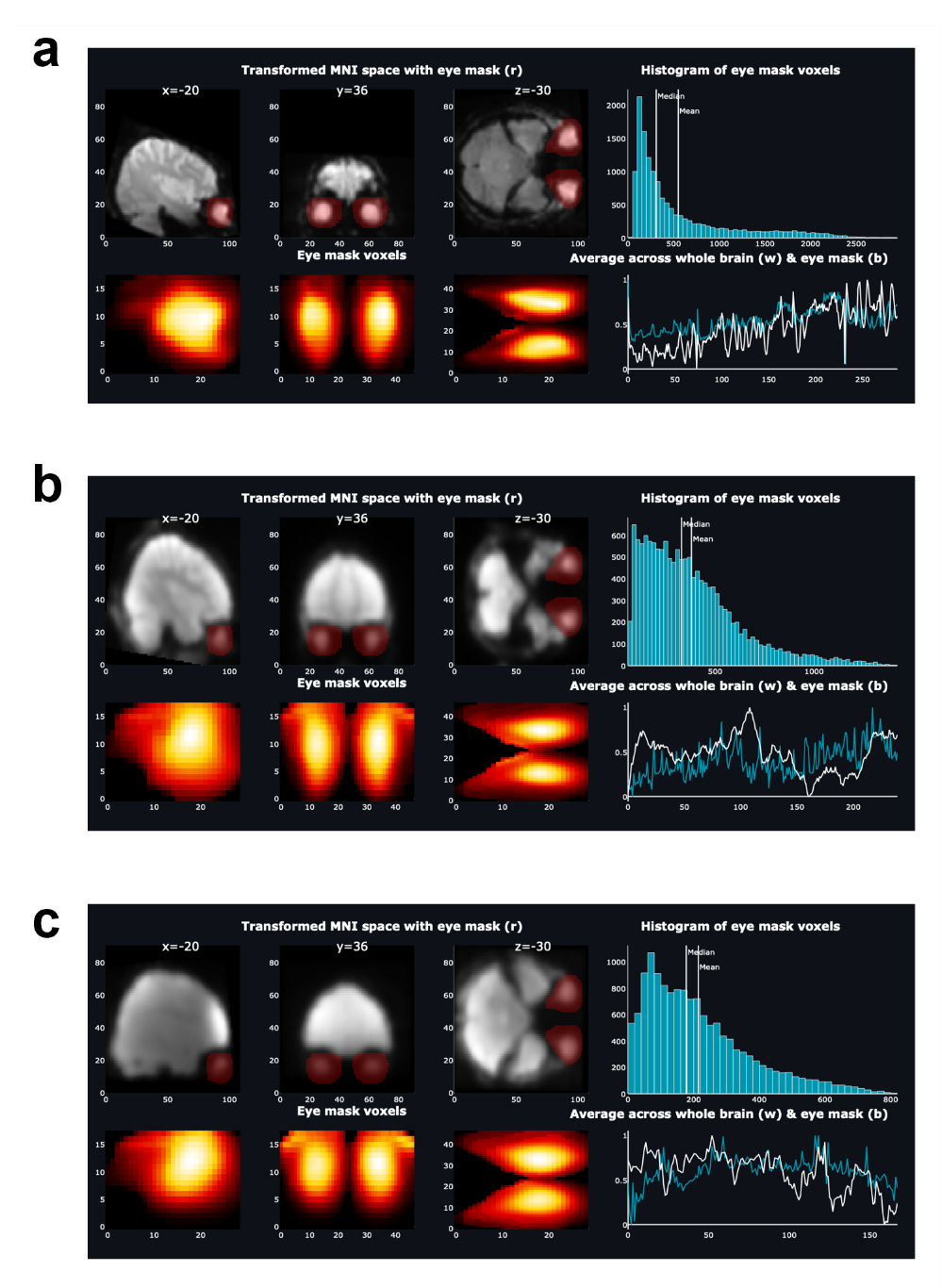

**Figure S2. Relationship between ground-truth eye-tracking data quality and model prediction accuracy in the Natural Viewing (NV) dataset.**

(a-b) Data from the *Despicable Me* (DM) stimulus. Panel (a) shows the association between the proportion of valid eye-tracking samples retained per participant and their DeepMReye prediction accuracy (Pearson’s *r*). Panel (b) shows the association between eye-tracking spatial precision—quantified as the root mean square (RMS) of sample-to-sample gaze displacement—and prediction accuracy.

(c-d) Parallel analyses for *The Present* (TP) stimulus. Panel (c) displays prediction accuracy as a function of the valid data proportion, and panel (d) shows accuracy as a function of RMS error. Shaded regions represent 95% confidence intervals.

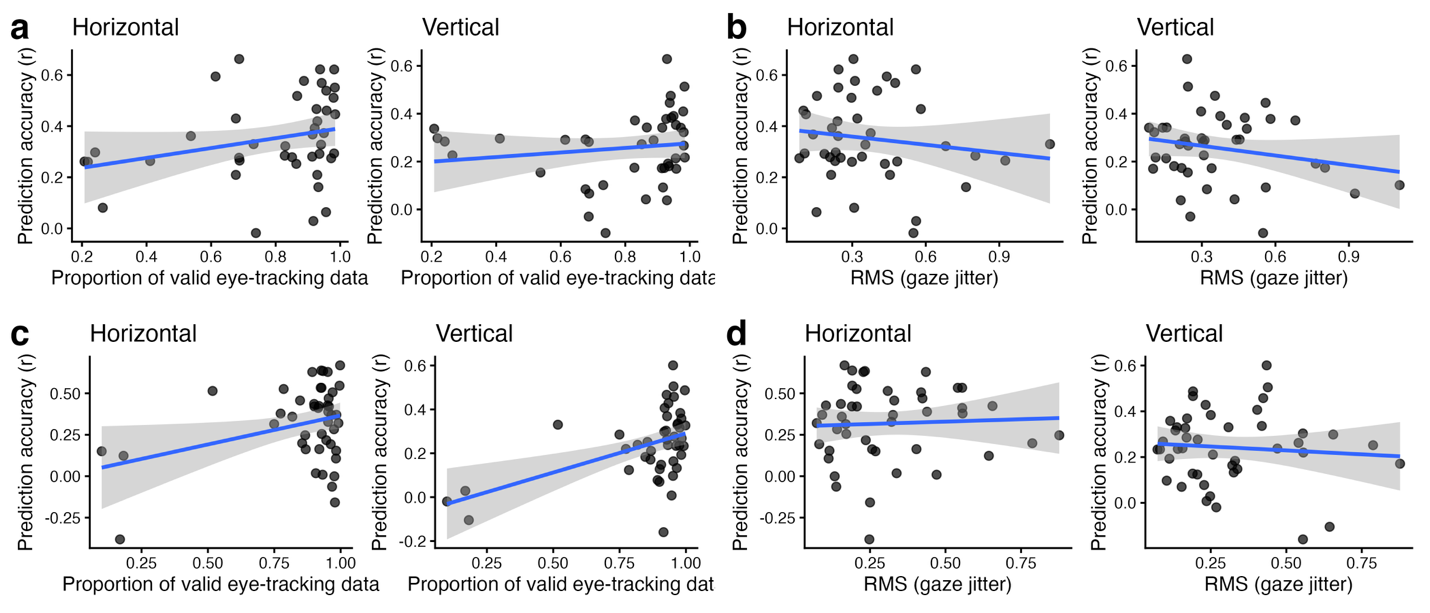

**Supplementary Figure S3. Relationship between the model's Predicted Error (PE) and gaze prediction performance in the Natural Viewing dataset.**

Model performance was evaluated using Pearson correlation (*r*) and Mean Absolute Error (MAE) across participants, stratified into low-PE and high-PE groups using a median split.

(a) Pearson correlation in the *Despicable Me* (DM) dataset for horizontal and vertical gaze components. Two sample t-tests revealed a significant difference for the vertical component, with higher values in the high-PE group.

(b) MAE in the DM dataset, showing no significant differences between PE groups.

(c) Pearson correlation in *The Present* (TP) dataset, showing no significant differences between groups for either gaze component.

(d) MAE in the TP dataset, demonstrating significantly higher spatial error in the high-PE group for both horizontal and vertical components.

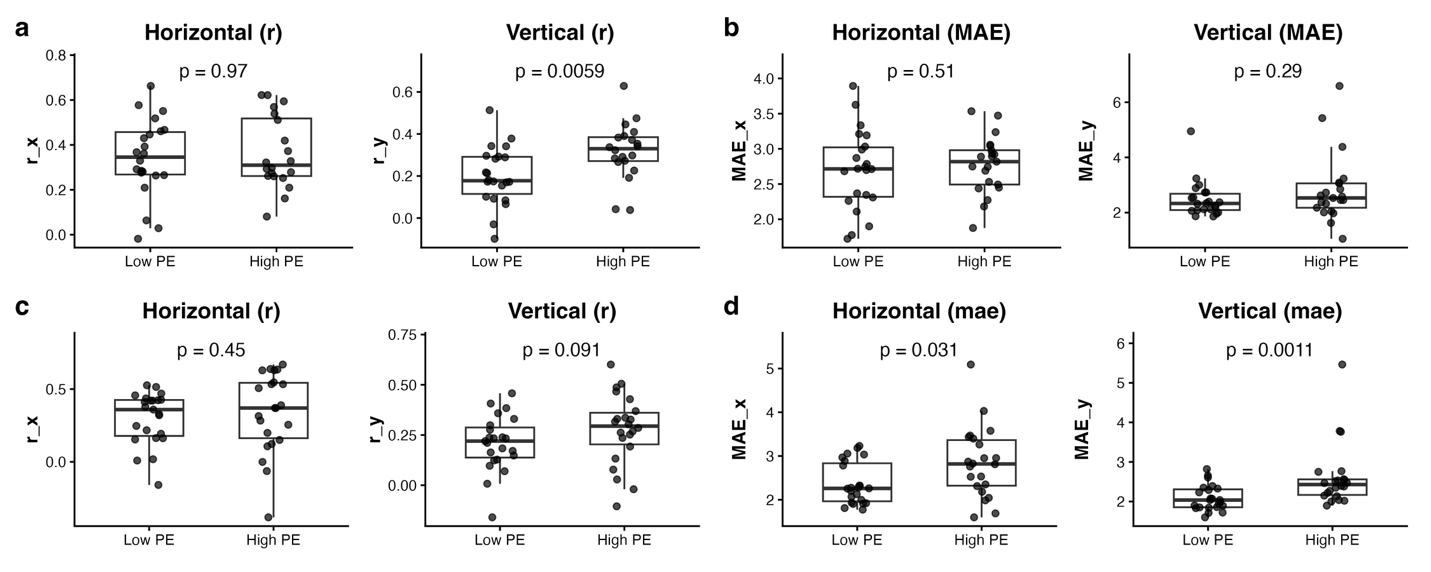
